# Predicting gene expression level in *E. coli* from mRNA sequence information

**DOI:** 10.1101/089102

**Authors:** Linlin Zhao, Nima Abedpour, Christopher Blum, Petra Kolkhof, Mathias Beller, Markus Kollmann, Emidio Capriotti

## Abstract

**Motivation:** The accurate characterization of the translational mechanism is crucial for enhancing our understanding of the relationship between genotype and phenotype. In particular, predicting the impact of the genetic variants on gene expression will allow to optimize specific pathways and functions for engineering new biological systems. In this context, the development of accurate methods for predicting translation efficiency from the nucleotide sequence is a key challenge in computational biology.

**Methods:** In this work we present *PGExpress*, a binary classifier to discriminate between mRNA sequences with low and high translation efficiency in *E*. *coli*. *PGExpress* algorithm takes as input 12 features corresponding to RNA folding and anti-Shine-Dalgarno hybridization free energies. The method was trained on a set of 1,772 sequence variants (WT-High) of 137 essential *E*. *coli* genes. For each gene, we considered 13 sequence variants of the first 33 nucleotides encoding for the same amino acids followed by the superfolder GFP. Each gene variant is represented sequence blocks that include the Ribosome Binding Site (RBS), the first 33 nucleotides of the coding region (C33), the remaining part of the coding region (CC), and their combinations.

**Results:** Our logistic regression-based tool (*PGExpress*) was trained using a 20-fold gene-based cross-validation procedure on the WT-High dataset. In this test *PGExpress* achieved an overall accuracy of 74%, a Matthews correlation coefficient 0.49 and an Area Under the Receiver Operating Characteristic Curve (AUC) of 0.81. Tested on 3 sets of sequences with different Ribosome Binding Sites, *PGExpress* reaches similar AUC. Finally, we validated our method by performing in-house experiments on five newly generated mRNA sequence variants. The predictions of the expression level of the new variants are in agreement with our experimental results in *E*. *coli*.

**Availability:** http://folding.biofold.org/pgexpress

## 1 Introduction

The ability to predict translation efficiency in bacteria is important to define the relation between genotype and phenotype, and to engineer new organisms optimized for producing biomaterials (Kyle, et al., 2009), fuels (Toone and de Winde, 2013) and natural products (Krivoruchko and Nielsen, 2015). The information to regulate the translation process is encoded in the mRNA nucleotide sequence. The preference for specific combinations of nucleotides in the coding region, which results in codon bias, has a strong effect on protein expression and formation (Li, et al., 2012; Mortimer, et al., 2014; Plotkin and Kudla, 2011; Pop, et al., 2014). Changes in the nucleotide sequence and codon usage can affect the mRNA folding process, which is a key determinant of protein expression. The ability of RNA strands to fold and form stable structures influences all the steps of the translation process: initiation, elongation, mRNA localization and turnover (Bentele, et al., 2013; Bonde, et al., 2016; Duval, et al., 2013; Goodman, et al., 2013; Mortimer, et al., 2014). The Shine-Dalgarno (SD) sequence encoded in the mRNA is another key factor for translation regulation. Indeed, when the SD sequence is located in untranslated regions (UTRs), it promotes the binding of ribosomes and accelerates translational initiation (Kozak, 2005; Shine and Dalgarno, 1974; Shultzaberger, et al., 2001). Contrarily, its presence in the coding region can reduce the translational elongation rate in bacteria (Li, et al., 2012). Thus, understanding bacterial translation on a mechanistic level will result in accurate predictions of protein expression from mRNA sequence (Gingold and Pilpel, 2011). In this work we considered the measure of translation efficiency, which provides a quantitative estimation of the of translation process, independent from the transcription. The translation efficiency is defined as the ratio of protein to mRNA abundance, which corresponds to the amount of protein produced by a single molecule of mRNA (Tuller, et al., 2010a; Tuller, et al., 2010b).

In the past, many studies and software tools have been developed for predicting protein expression based on mRNA sequence. Tools to tailor the untranslated region (UTR) to achieve a desired protein expression level were also introduced (Na and Lee, 2010; Reeve, et al., 2014; Rodrigo and Jaramillo, 2014; Seo, et al., 2014). The RBS calculator (Salis, 2011), UTR designer (Seo, et al., 2014), and RBS designer (Na and Lee, 2010) are statistic methods based on thermodynamic models. They calculate the folding free energies for key molecular interactions to provide an estimation of the translation efficiency. In general, the predictions from these methods show high correlation with experimental data. Recently Bonde and colleagues (Bonde, et al., 2016) studied the relationship between SD sequences and protein expression by measuring expression levels of ~3,000 UTRs in the presence of different SD variants. Their empirical method (EMOPEC) outperformed the standard thermodynamic models. Considering only the UTR regions, the available tools limit our understanding of the general picture of translational mechanism and our ability to engineer the whole mRNA molecule. Goodman and colleagues (Goodman, et al., 2013) measured the expression level of more than 14,000 synthetic gene variants in *E. coli* to quantify the effects of N-terminus codons as well as different combinations of promoter and ribosomal binding sites (RBSs). They found that rare codons in the N-terminus increased the stability of the RNA structure resulting in decreased gene expression level. The gene variants tested by Kosuri and co-workers (Goodman, et al., 2013) included variations in both UTR and coding sequences, which made the data suitable for investigating the effects from coding sequences as well. We make use of their data to capture regulatory factors from both the UTR and coding region of the mRNA molecule.

For estimating the contributions of different RNA regions on gene expression, we represented the sequences by global and local free energy features to find the main determinants of the translation efficiency. Since mRNA structure impacts each step of translation (Kozak, 2005; Mortimer, et al., 2014), it represents one the most important features to consider. The folding free energy is a classical measure to describe the RNA structure. Many tools to predict RNA structure implement thermo-dynamic-based dynamic programming algorithms (Capriotti and Marti-Renom, 2008). Many studies showed that different regions of mRNA preserve specific structural preferences (Kudla, et al., 2009; Mortimer, et al., 2014). Kudla and colleagues found that the predicted folding free energy of the first ~40 nucleotides of the mRNA has a significant correlation with the GFP protein abundance (Kudla, et al., 2009). In a recent study, it was observed that structures at the end of 5′ UTR and the beginning of 3′UTR are well conserved and the coding region is more structured than UTRs (Mortimer, et al., 2014). Thus, the free energy associated to the formation of local structures is also an important predictive feature. In addition, since the SD sequence shows different regulating effects, we also computed the hybridization folding energy (also referred as binding energy) between the anti-SD sequence and different regions of the mRNA. The folding and hybridization free energies were combined to represent the translational features of the mRNA.

In this work we present *PGExpress* (*Predicting Gene Expression*), a new logistic regression-based algorithm to predict translation efficiency of mRNA sequences and designing new gene variants for experimental testing. *PGExpress* is a binary classifier that discriminates between nucleotide sequences with low and high translation efficiency. Our method relies on the calculation of the minimum free energy of folding as representations of the local and global mRNA structures and the minimum free energy of hybridization between anti-SD sequence and mRNA, which corresponds to the binding affinity of the ribosome with different strands of mRNA. The performance of *PGExpress* has been tested on previously published datasets and new experimental data generated in-house.

## 2 Methods

### 2.1 Datasets

The data used in this work consists of protein expression and/or translation efficiency measures of genes and their variants in *E. coli.* The data were collected both from the literature (Goodman, et al., 2013; Taniguchi, et al., 2010) and experimental tests in our lab. The data from Kosuri and collaborators (Kosuri-All) is a collection of protein expression and translation efficiency measures from ~ 14,000 gene variants (Goodman, et al., 2013). Each variant is a combination of the Promoter with high and low strength (High, Low), the Ribosome Binding Site (Wild-Type, Weak, Mid and Strong RBSs) and the first 33 nucleotides of the coding region (C33) of 137 essential *E. coli* genes followed by the superfolder GFP (sfGFP) coding sequence (see Supplementary Materials, section Experimental data). From the Kosuri-All dataset we extracted five subsets (WT-High, WT-Low, Weak-High, Mid-High, Strong-High) with sequence variants composed by four Ribosome Binding Sites (RBS) and two Promoters. The main dataset (WT-High), which has been used for training and testing our method, collects the expression measures of 1,722 sequences formed by the High promoter, the Wild-Type RBSs and 13 variants (including wild-type) of the C33 region of each gene (see Supplementary File 1). The Weak-High, Mid-High and Strong-High subsets, which have been used only for the testing phase, differ from the WT-High for the sequence of the Ribosome Binding Site, which have Weak, Mid and Strong binding affinities respectively (see Supplementary Files 2, 3 and 4 respectively). The WT-Low and WT-High differ for the sequence of the promoter regions, which have low and high strength respectively. The WT-Low dataset has been used only in the preliminary analysis of the data (see Supplementary File 5). All gene variants are classified in two classes according to their translation efficiency. The median value of the translations efficiency in WT-High (2355.5) was used as classification threshold. To test the performance of *PGExpress*, we measured in our lab the protein expression level of five randomly selected variants from the Kosuri-All dataset (Exp-Set). We used the Exp-Set to check the agreement between the data in Kosuri-All and our measures. Additionally, we generated a validation set, namely Exp-Mut, which is composed of new variants derived from the five sequences in Exp-Set. The sequences of the ten gene variants are reported in Table S1.

Finally, we estimated the ability of our algorithm to predict the expression level of the full gene comparing the translation efficiency from Kosuri’s study with the single-cell data from a recent work in which the protein expression of wild-type genes was measured (Taniguchi et al., 2010). For this comparison, we selected a subset of 44 common genes (Xie-Comm), for which the expression level was reported in High-WT and experimentally determined by Xie and collaborators (see Supplementary File 6). Finally, we focused on a subset of 29 genes on which both studies are in agreement (Xie-Agree). In other words, Xie-Agree dataset is composed by the subset of genes for which the expression level from Xie-Agree (Taniguchi, et al., 2010) and the translation efficiency from Kosuri-All (Goodman, et al., 2013) are either higher or lower with respect to their median values. Since the common genes analyzed in both studies differ only for the composition of the C-terminal region (CC), we assume that our filtering procedure results in the selection of the group of genes for which the contribution of the C-terminal region of the coding sequence has little effect on the translation process.

All the datasets used in this work are provided as supplementary files and a summary of their composition is reported in Table S2.

### 2.2 Algorithm description

Here we present a binary classifier (*PGExpress*) to predict the gene translation efficiency from sequence information. *PGExpress* is based on logistic-regression algorithm that takes in input a 12-elements vector composed by six RNA folding and six anti Shine-Dalgarno (SD) hybridization free energies. In detail, each gene variant is divided in three sequence blocks: the Ribosome Binding Site (RBS), which consists on average of 20 nucleotides preceding the coding sequence, the first 33 nucleotides of the coding region (C33) and the remaining part of the coding sequence starting from nucleotide 34 (CC). Thus, each gene is represented by six sequence fragments including the three blocks previously defined (RBS, C33 and CC), and the combinations of RBS with C33 (RBS+C33), C33 with CC (CDS) and RBS with the whole coding sequence (RBS+CDS). For each block we calculated the RNA folding and the anti-Shine-Dalgarao (anti-SD) hybridization free energies using respectively *RNAfold* and *RNAduplex* tools from the ViennaRNA package (Lorenz, et al., 2011), which automatically replace Thymine (T) with Uracil (U). We used an 8-nucleotides anti Shine-Dalgarno sequence (CCTCCTTA) as reported by Kosuri and coworkers (Goodman, et al., 2013). Both free energies have been rescaled to a temperature of 30 °C, which is the temperature at which the experiment in the Kosuri study was carried out. *PGExpress* uses a softmax function to calculate the probabilistic score (P) returned as output. If P>0.5, the gene variant is predicted to have a high translation efficiency (Trans>2355.5). A representation of *PGExpress* and its 12 input features is provided in Fig. 1.

**Fig. 1.**
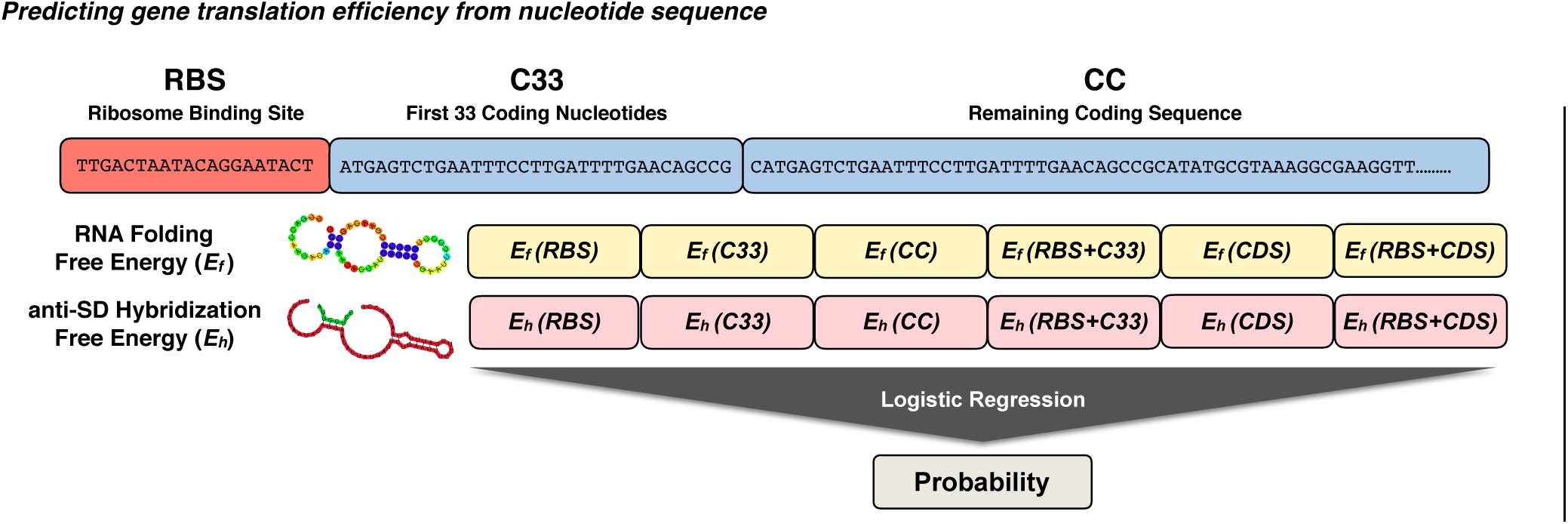
Representation of the *PGExpress* algorithm. *PGExpress* input is a 12-elements vector composed by six RNA folding (*E*_*f*_) and six anti-Shine-Dalgarno hybridization (*E*_*h*_) free energies. Each sequence is divided in 3 blocks: the Ribosome Binding Site (RBS), the first 33 nucleotides of me coding sequence (C33) and the remaining part of the coding region starting from nucleotide 34 (CC). The whole coding sequence (CDS) is obtained joining C33 and CC.

### 2.3 Feature analysis

To estimate the discriminative power of each feature, we compared the distributions of the RNA folding and anti-SD hybridization free energies of the five sequence blocks (RBS, C33, RBS+C33, CDS and RBS+CDS) on the subset of variants with high (Trans>2355.5) and low (Trans ≤2355.5) translation efficiencies in the WT-High dataset. In this analysis we did not consider the C-terminal region of the coding sequence (CC) because it corresponds to the sfGFP for all the variants in the Kosuri-All dataset. The comparison between the two distributions is performed calculating the Kolmogorov-Smirnov distance and the associated p-value. Furthermore, we compared the performance of our best approach (*PGExpress*) against five methods including different combinations of the 12 input features. These methods are:

- **BFolding**: most discriminative RNA folding free energy
- **BBinding**: most discriminative anti-SD hybridization free energy
- **Folding6:** RNA folding free energies of the six blocks
- **Binding6:** anti-SD hybridization free energies of the six blocks.
- **BFoldBind:** most discriminative RNA folding and anti-SD hybridization free energies.

### 2.4 Algorithm optimization

*PGExpress* is based on a logistic-regression algorithm (LogisticRegression) implemented in the *scikit-learn* package (Pedregosa, et al., 2011). It has been optimized considering different tolerance values (0.1, 0.05, 0.01, 0.005 and 0.001). The *scikit-learn* LogisticRegression class was run using the L1 regularization method as penalty function and the defaults values for all the remaining options.

### 2.5 Training and testing

To estimate the performance of *PGExpress* and the alternative methods, we performed several tests. First, we tested *PGExpress* using a gene-based 20-fold cross-validation approach on the WT-High dataset to keep all the variants belonging to the same gene in the same subset. For each test we calculated the performance using the evaluation measures defined in Supplementary Materials. The reported scores represent the average values obtained over five 20-fold cross-validation tests. The results obtained on the Kosuri-All (Weak-High, Mid-High and Strong-High), Experimental (Exp-Set, Exp-Mut) and Xie (Xie-Comm, Xie-Agree) datasets were calculated removing from the training set all the data related to the genes present in the testing set. This procedure reduces the overfitting due to the presence of data from sequences with high similarity both in training and testing sets. To check for this source of bias, we also performed the all-against-all global alignments (1,558,513) among the RBS+C33 regions of all the gene variants. The global alignments of the nucleotide sequences were calculated using the *align0* algorithm from the *fasta2.0* package (Myers and Miller, 1988).

### 2.6 Engineering new sequences

For validating our algorithm, we generated new sequences selecting the subset of gene variants which showed by mutation either higher or lower translation efficiency with respect to the median value of the High-WT dataset. For calibrating our experimental expression measures and compare them with the data reported by Kosuri and colleagues (Goodman, et al., 2013), we first measured the expression level of five randomly selected gene variants (Exp-Set). In the next step, we generated five new sequences not included in the Kosuri-All dataset mutating at most one nucleotide in RBS or three codons in coding region. Finally, we randomly selected a set of five gene variants (Exp-Mut), four of which change their expression level either from High to Low *(dapB* and *lpxK)* or Low to High *(lgt* and *zipA)* and one case *(murF)* where the expression level remains in the same class. The sequences of the ten tested gene variants are reported in Table S1.

### 2.7 Experimental expression measure

DNA sequences consisting of promoter, ribosomal binding site (RBS), and 33 coding nucleotides (including ATG start site) of five different genes were synthesized (Genscript, Piscataway, USA) with flanking AscI and NdeI restriction sites. The DNA fragments were excised from the shuttle vector and directionally cloned into the pJ251-GERC vector obtained from Addgene (Kosuri, et al., 2013). A unique EcoRI restriction site was engineered in between the 5′ region of the AscI site and the respective promoter sequence. Using the EcoRI site we identified the positive clones. Final gene variants were verified via Sanger sequencing. The correct variants were transformed in MG165 *E. coli* cells and starter cultures were grown over night at 37 °C. The next day cultures were diluted 1:1000 in 100 μL LB medium in optical quality black walled 96-well plates (PerkinElmer, Waltham, MA, USA) in quadruplicate and overlayed with 40 μL mineral oil. Bacteria were grown at 30 °C. Bacterial growth was followed by measuring the optical density at 600 nm (OD600) as proxy. The different combination of promoter, RBS, and coding region regulate the expression levels of the superfolder green fluorescent protein (sfGFP). Expression of the red fluorescent protein (mCherry) was controlled by a constitutive promoter (PLtetO-1) shared by all gene variants (Kosuri, et al., 2013). sfGFP and mCherry fluorescence levels were measured with a monochromator equipped BioTek Synergy Mx (BioTek, Winooski, USA) plate reader. Every five minutes a fluorescence measurement was performed.

## 3 Results

### 3.1 Classification and input features

The selection of the data from Kosuri and co-workers allowed us to develop a machine learning method *(PGExpress)* that classifies the gene variants as having low (Trans≤2355.5) and high (Trans>2355.5) translation efficiency based on the sequence information. Before performing our tests, we analyzed the Kosuri-All dataset and focused on the gene variants in the WT-High subset. This set is composed of sequences with promoter with high binding affinity (BBaJ23100) and wild-type RBSs (Ribosome Binding Sites). The choice of WT-High dataset is supported by the observation that the correlation between the level of protein and RNA expression is higher than in WT-Low dataset (Fig. S1). Indeed, the correlation coefficients are 0.72 and 0.51 in WT-High and WT-Low sets, respectively. Thus, we selected the WT-High subset as a main reference set for this work. We divided the WT-High dataset in two subsets of gene variants with low and high translation efficiency with respect to the median value. In addition, the WT-High dataset was used to estimate the predictive power of our machine learning approach. To avoid the overestimation of the performances we performed a gene-based 20-fold cross-validation test. Keeping the variants from the same gene in the same subsets, we excluded the presence of sequences with high level of identity both in training and testing. Thus, we calculated the distribution of the percentage of identity (PID) between the first two blocks (RBS+C33) of the different gene variants. The Fig. S2 shows that only ~4% of the cases the PID achieved a value between 50% and 60%. To estimate the predictive power of the input features used in *PGExpress*, we performed the Kolmogorov-Smirnov (KS) test between their distributions in the subset of variants with low and high translation efficiency. The Tables S3 and S4 report the KS distance, the associated p-value and other statistical measures for the distributions of RNA folding and anti-Shine-Dalgarno (anti-SD) hybridization free energies. This analysis revealed that overall the free energies of the RBS+C33 sequences result in the highest divergence between of the distributions of low and high expressed variants. Indeed the RBS+C33 score distributions correspond to the lowest RNA folding (Tables S3) and the second lowest anti-SD (Tables S4) p-values. This observation is confirmed by the boxplot in Fig. S3 which shows the distribution of RNA folding (panel A) and anti-SD hybridization (panel B) free energies of the gene variants with low (black) and high (white) translation efficiency.

### 3.2 Performance of different methods

In a second step, we calculated the real performance of *PGExpress* and five alternative methods, which included a reduced number of features. The input features for the BFolding, BBinding, Folding6, Binding6 and BFoldBind were described in the section Feature Analysis. In Table 1 we tested the performance of the six methods on the WT-High dataset using the gene-based 20-fold cross-validation procedure. The results revealed that the RNA folding free energy corresponding to the RBS+C33 portion of the variants is the most informative feature. Indeed the BFolding method reached an overall accuracy (ACC) of 0.68 a Matthews Correlation Coefficient (MC) of 0.37 and Area Under the Receiver Operating Characteristic Curve (AUC) of 0.74.

**Table 1.** Performance of the methods using alternative input features. ACC, MC, AUC and other evaluation measures are defined in Supplementary Materials. N is the number of input features. The input features of BFolding, BBinding, Folding6. Binding6. BFoldBind and *PGExpress* are defined in the section Features analysis.

| Method | ACC | TNR | NPV | TPR | PPV | MC | AUC | N |
| --- | --- | --- | --- | --- | --- | --- | --- | --- |
| BFolding | 0.68 | 0.63 | 0.70 | 0.73 | 0.67 | 0.37 | 0.74 | 1 |
| BBinding | 0.54 | 0.50 | 0.55 | 0.59 | 0.54 | 0.09 | 0.55 | 1 |
| Folding6 | 0.69 | 0.64 | 0.71 | 0.74 | 0.67 | 0.38 | 0.74 | 6 |
| Binding6 | 0.54 | 0.53 | 0.54 | 0.55 | 0.54 | 0.07 | 0.56 | 6 |
| BFoldBind | 0.72 | 0.70 | 0.73 | 0.74 | 0.71 | 0.44 | 0.79 | 2 |
| *PGExpress* | 0.74 | 0.72 | 0.75 | 0.76 | 0.73 | 0.49 | 0.81 | 12 |

The discriminative power of the anti-Shine-Dalgarno (anti-SD) binding free energy is much lower. This is evident by measuring the performance of the BBinding method that achieved an ACC of 0.54 and MC of 0.08 and AUC of 0.55. The analysis of the results of the Folding6 and Binding6 methods, which include six features of the same type free energy, do not show any substantial increase in the performances with respect to the BFolding and BBinding methods. The first improvement in the performance is obtained combining the RNA folding and anti-SD hybridization free energies of the RBS+C33 sequence. Indeed the BFoldBind method, which takes in input only two features, reached an ACC of 0.72 a MC of 0.44 and AUC of 0.79. In *PGExpress* we merged the six RNA folding and six anti-SD hybridization free energies. The results in Table 1 show that *PGExpress* achieved and ACC of 0.74, a MC of 0.49 and AUC 0.81 improving the Matthews correlation coefficient of 0.05 and the AUC of 0.02 with respect to BFoldBind. The Receiver Operating Characteristic (ROC) curves for all methods are plotted in Fig. 2A. The little improvement resulting from the usage of 12 features is due to absence in the training set of variants in the 3’ region of gene which starts from nucleotide 34 (CC). Indeed in all the experiments reported by Kosuri and coworkers the CC block corresponds to the superfolder GFP coding sequence.

**Fig. 2.**
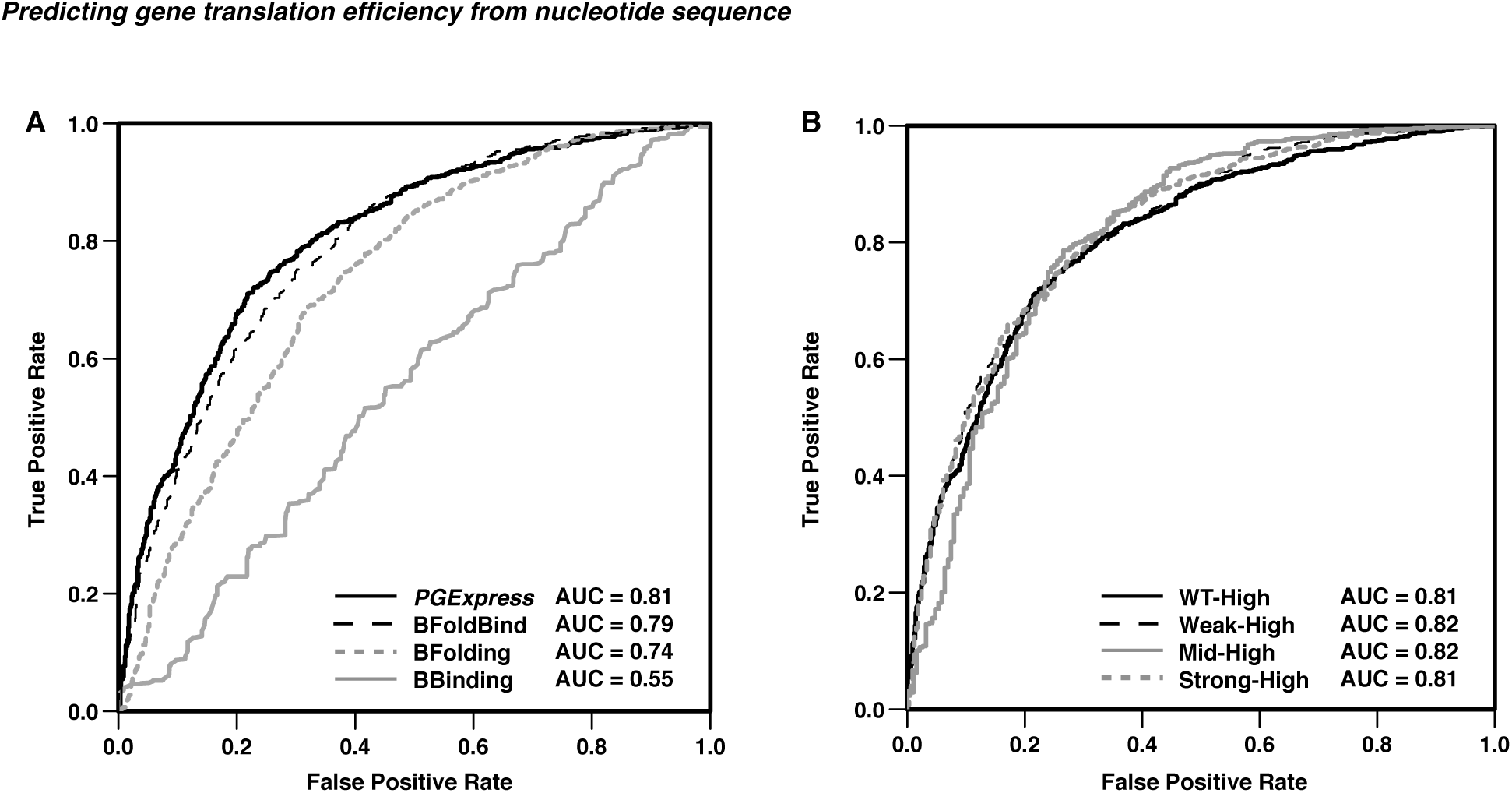
ROC curve of the predictor. (A) ROC curves *PGExpress* and alternative metiiods with reduced input features on the WT-High dataset. (B) ROC curves of *PGExpress* on WT-High. Weak-High, Mid-High and Strong-High datasets.

Although the Kosuri-All dataset presents this limitation, for generalization purpose, we decided to include the CC energy scores in the input features of *PGExpress*. Our algorithm was optimized testing different values of tolerance for the logistic regression algorithm. The results in Table S5 do not show strong differences among the tested tolerance values, thus we select a tolerance equal to 0.05 as parameter for the training.

### 3.3 Performance on the Kosuri-All subsets

In the next test we focused on the performance of *PGExpress* on three datasets (Weak-High, Mid-High and Strong-High), which contain gene variants with the same 33 starting nucleotides in the coding regions (C33) but three different RBSs (Ribosome Binding Sites). Analyzing the three new datasets, we observed that the distribution of the translation efficiency (Trans) in Weak-High and WT-High are similar while Mid-High and Strong-High are strongly unbalanced toward high translation efficiency values (see Fig. S4). Thus, comparing the performance on WT-High with those on the three new datasets, we observed that *PGExpress* achieved slightly lower overall accuracy (ACC) and Matthews correlation coefficient (MC) on the Weak-High dataset and, due to the dataset unbalance, higher ACC and lower MC on Mid-High and Strong-High (Table 2). Nevertheless, the Area Under the ROC Curves for all the datasets are similar (Fig. 2B).

**Table 2.** Performance of the PGExpress on the Kosuri-All subsets. ACC, MC, AUC and other evaluation measures are defined in Supplementary Materials. %High is the percentage of gene variants with high translation efficiency.

| Dataset | ACC | TNR | NPV | TPR | PPV | MC | AUC | %High |
| --- | --- | --- | --- | --- | --- | --- | --- | --- |
| WT-High | 0.74 | 0.72 | 0.75 | 0.76 | 0.73 | 0.49 | 0.81 | 50 |
| Weak-High | 0.73 | 0.59 | 0.78 | 0.85 | 0.70 | 0.46 | 0.82 | 53 |
| Mid-High | 0.83 | 0.12 | 0.84 | 1.00 | 0.83 | 0.28 | 0.82 | 81 |
| Strong-High | 0.90 | 0.13 | 0.74 | 0.99 | 0.91 | 0.29 | 0.81 | 89 |

### 3.4 Selecting high-quality predictions

To better characterize the performance of *PGExpress*, we scored our method filtering-out the less reliable training data and predictions in WT-High dataset. First we assume that gene variants with translation efficiency near the median (2355.5) constitute the noisy part of the dataset. Thus we re-scaled all the translation efficiency measures calculating Logarithm Ratio (LS) with respect to the median value (see Evaluation Measures section in Supplementary Materials) and filtered-out the data below a given threshold. The performance of *PGExpress* after removing the data close to the median value are reported in Fig. 3A and Table S6. We observed that removing 42% of the gene variants with translation efficiency higher than 1177.7 and lower than 4711 *(LR<1), PGExpress* reached an overall accuracy of 0.81 and an AUC of 0.88. Similar analysis was performed filtering-out less reliable predictions. We used the probabilistic output (P) of our machine learning method to calculate the Reliability Index (R1) as absolute value of the difference P-0.5, re-scaled between 0 and 10. The results in Fig. 3B and Table S7 shows that removing the predictions with RI<3, *PGExpress* achieved an accuracy of 0.81 and an AUC of 0.85 on 63% of the WT-High dataset.

**Fig. 3.**
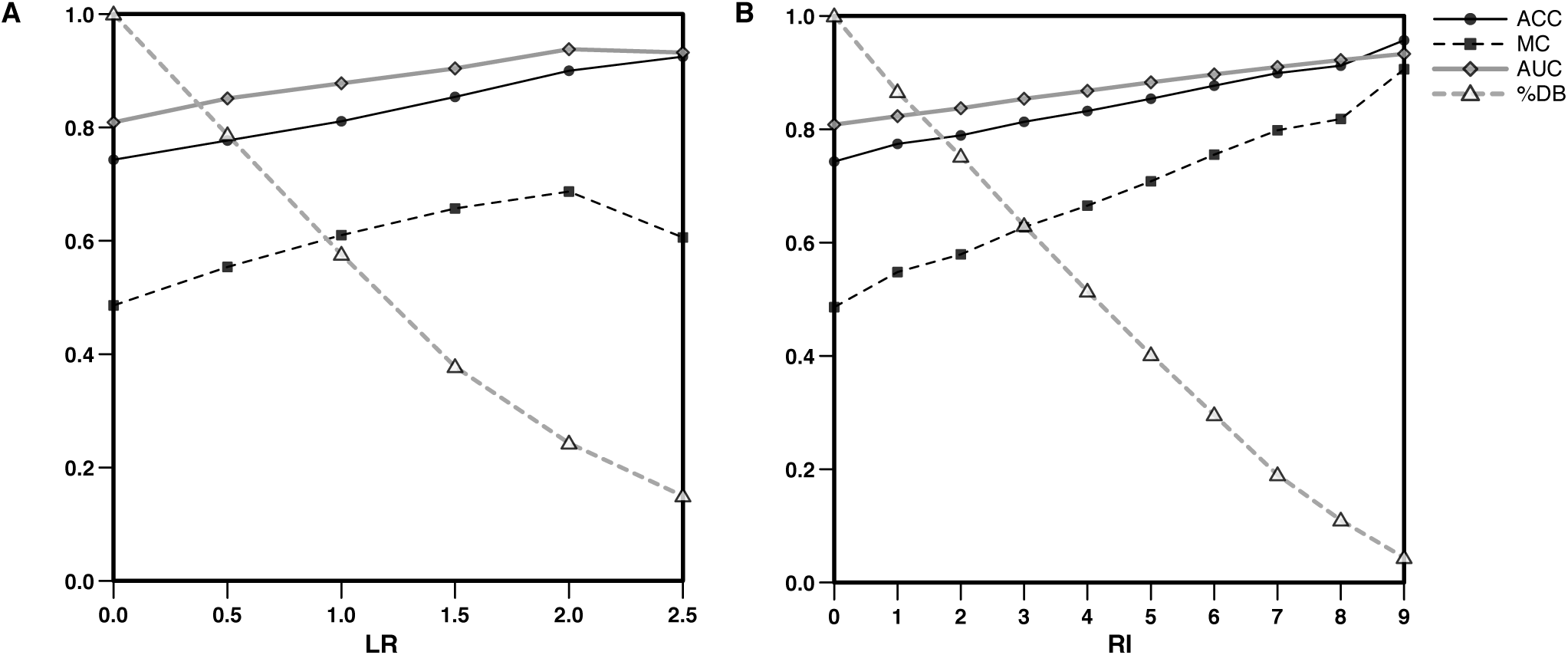
Performance of the method as a function of the Logarithm Ratio (LR) and Reliability Index (RI). LR, RI, ACC, MC and AUC are defined in Supplementary Materials. %DBs the percentage of the dataset after filtering out less reliable training data (panel A) or predictions (panel B).

### 3.5 Test on our experimental dataset

To test the ability of *PGExpress* to predict the translation efficiency we performed in-house experiments with five gene variants each in the Exp-Set and Exp-Mut datasets (see methods section) and measured the protein expression using the protocol introduced by Kosuri and coworkers (Goodman *et al*., 2013). In Fig. S5 we plotted the measures of the fluorescence associated to each gene variant normalized by the maximum level of OD600. To make a fair comparison between our results and those reported by Kosuri and collaborators, we used the median value of the protein expression level in Kosuri data as threshold for discriminating between low and high expressed gene variants. Thus, we compared the maximum value of the re-scaled fluorescence (Table S8 and Fig. S5) obtained in our experiment with the median protein expression level in the WT-High dataset (2998.1).

According to this assumption, we verified that for four gene variants over five (Exp-Set), our experiments match those performed by Kosuri and colleagues (Table 3). The only difference is observed for a variant of the *lgt* gene (lgt-23), which is classified to have high protein expression and translation efficiency in the Kosuri-All dataset, whereas our experiments revealed a low protein expression. Nevertheless the prediction of *PGExpress* agrees with the results reported in Kosuri-All dataset. Finally we evaluate the accuracy of *PGExpress* predictions on the Exp-Mut dataset, verifying that our predictions are correct for all the new gene variants. A dubious prediction is represented by the variants lpkX-Mut which is predicted to have low expression level (P=0.19) and, our in-house measure of protein expression (2996), is only few digits below the threshold (2998.1).

**Table 3.** Prediction of the expression level for the gene variants (ID) in the Exp-Set and Exp-Mut datasets. Kosun-All: translation efficiency from Kosun’s dataset Expenment: protein expression levels from our in-house experiments. High and Low are referred to the median values of the translation efficiency (2355.5) for Kosun-All data and the expression level (2998) for the Experiment data. Prediction: *PGExpress* predicted classes. Output: Probabilistic output of *PGExpress* defined m the section “Algorithm Description”. ^*^Our experimental measure for the lst-23 gene variant is in disagreement with data from Kosuri dataset. The sequences of all variants are reported m Table S1.

| Dataset | ID | Kosuri-All | Experiment | Prediction | Output |
| --- | --- | --- | --- | --- | --- |
| Exp-Set | dapB-28 | High | High | High | 0.73 |
|  | lgt-23* | High | Low | High | 0.55 |
|  | lpxK-30 | High | High | High | 0.90 |
|  | murF-21 | Low | Low | Low | 0.41 |
|  | zipA-23 | Low | Low | Low | 0.22 |
| Mut-Set | dapB-Mut | - | Low | Low | 0.23 |
|  | lgt-Mut | - | High | High | 0.78 |
|  | lpxK-Mut | - | Low | Low | 0.38 |
|  | murF-Mut | - | Low | Low | 0.19 |
|  | zipA-Mut | - | High | High | 0.88 |

### 3.6 Towards predictions from complete gene sequences

In the last part of this work we estimated the performance of *PGExpress* in predicting the translation efficiency of the full-length wild-type gene. For this analysis we compared the data from Kosuri and colleagues, where the 3′ region of the gene starting from the nucleotide 34 (CC) was replaced by sfGFP, with the experimental results from Xie and coworkers in which the full sequence of the gene was considered (REF). The direct comparison of the two studies is a challenging task, in which the differences in the experimental setting can lead to contradictory results. Thus, we extracted a set of 44 common genes (XieComm) and divided it in two groups. The Xie-Agree is a reliable set of 29 genes in which we assumed that both studies are in agreement because they are performed in comparable experimental conditions and the contribution of the 3’ region is not significant. The second subset of 15 genes (XieDiff) for which the predictions can be strongly affected by the experimental conditions and/or the contribution of the 3’ region. The results in Table 4 show that the performance of *PGExpress* on the XieComm dataset is close to random with ~55% overall accuracy and 0.51 AUC. After the filtering procedure, that removed the genes with contradictory expression levels, *PGExpress* achieved an overall accuracy of ~81%, Matthews correlation coefficient of 0.52 and AUC equal to 0.83.

**Table 4.** Performance of PGExpress on the Xie subsets. N is the number of genes in each set. ACC, MC, AUC and other accuracy measures are defined in Supplementary Materials.

| Dataset | ACC | TNR | NPV | TPR | PPV | MC | AUC | N |
| --- | --- | --- | --- | --- | --- | --- | --- | --- |
| XieComm | 0.55 | 0.42 | 0.28 | 0.60 | 0.74 | 0.02 | 0.51 | 44 |
| XieAgree | 0.81 | 0.71 | 0.58 | 0.84 | 0.90 | 0.52 | 0.83 | 29 |
| XieDiff | 0.07 | 0.00 | 0.00 | 0.10 | 0.17 | -0.87 | 0.00 | 15 |

## 4 Discussion

In this work we presented *PGExpress*, a logistic regression-based method for predicting translation efficiency of mRNA from a set of free energy features. The method uses the folding free energies of six sequence blocks which represent the local and global mRNA folding structures. The six blocks include RBS, C33, CC sequence and their combinations. Among them the folding energy the block of RBS+C33 provides the most informative feature. This is in agreement with previous findings that the folding structure around starting codon has a strong effect on translation. By adding folding free energies from other blocks, the prediction accuracy slightly increased. This might indicate that, although other regions of the gene have an impact on translation, the structure of the 5’ region constitutes the main contribution to the translation rate. For instance, the presence of a folded SD sequence near a starting codon might slow down the translation process reducing the probability of the ribosomes to bind or elongate. Accordingly, the minimum hybridization free energies were used to represent the effect of the SD sequence calculating the hybridization energy between the mRNA and the anti-SD sequence. Although the minimum hybridization free energy itself shows a weak correlation with the translation efficiency, the interplay between all folding and hybridization free energies allowed to improve the performance of our predictor. This indicates that the mRNA structures and SD sequences regulate translation in a cooperating manner. In addition, we test the sensitivity of *PGExpress* to small changes in the nucleotide sequences. For this purpose we measured the expression level of five gene variants that differ in few nucleotides from the original sequences from Kosuri-All dataset. Our analysis show that *PGExpress* is able to correctly predict the expression level of all new variants, most of which (4/5) resulted in an opposite expression level with respect to the original sequence.

Strikingly, is the case of the *dapB* variant which achieved ~ 10-fold lower expression with only 2 synonymous mutations (see Tables S1 and S8). This observation confirms the robustness of our method, which supports its practical application in biotechnology. Compared with other methods that are merely focusing on the effects of RBSs, we integrated the main effecting factors from the perspective of whole sequence, which enabled us to predict translation efficiency accurately and to engineer new sequences at the whole sequence level.

We also tested *PGExpress* on the dataset released by Xie and collaborators (Taniguchi, et al., 2010), which measured the expression level of the full sequence of each gene. By considering a subset of 29 full genes *PGExpress* achieved ~81% overall accuracy indicating that, for this subset of genes (XieAgree), the features extracted from the 5’ region of gene allowed to reach a good level of generalization. Further investigations are needed to understand the poor performance obtained on the remaining subset of genes (XieDiff). This result can be due to the inconsistency among the two experimental setting and/or the stronger contribution of the 3’ region to the expression level of these genes. In the latter case, the limited information available in Kosuri dataset, which do not includes full coding region of the genes, confines our ability to investigate further the translational mechanisms.

The binding and elongation in the translation process, which involves the RBS recognition and the sliding of the ribosomal complex along the whole mRNA sequence (Kudla, et al., 2009; Li, et al., 2012; Plotkin and Kudla, 2011), suggests that position specific features can provide a more realistic model for the translational mechanism. However the scale and composition of the dataset limit the application of more complex machine-learning methods based on position-specific features. Future directions of our work will include the analysis of new features to improve the prediction of the translation efficiency of wild-type genes in *E. coli*, and the development of tools for identifying key nucleotides to control protein expressions. We believe that our *in-silico* approach can have strong impact on biotechnological applications reducing the experimental effort to engineer optimized organisms.

## Acknowledgements

E.C. acknowledges Genifx - Genome Informatics Service at the University of Alabama at Birmingham, AL (USA) for computational resources. We acknowledge Sriram Kosuri, George Church and collaborators for sharing their experimental data.

